# 2Danalysis: A toolbox for analysis of lipid membranes and biopolymers in two-dimensional space

**DOI:** 10.1101/2025.02.27.640563

**Authors:** Ricardo X. Ramirez, Antonio M. Bosch, Rubén Pérez, Horacio V. Guzman, Viviana Monje

## Abstract

Molecular simulations expand our ability to learn about the interplay of biomolecules. Biological membranes, composed of diverse lipids with varying physicochemical properties, are highly dynamic environments involved in cellular functions. Proteins, nucleic acids, glycans and bio-compatible polymers are the machinery of cellular processes both in the cytosol and at the lipid membrane interface. Lipid species directly modulate membrane properties, and affect the interaction and function of other biomolecules. Natural molecular diffusion results in changes of local lipid distribution, affecting the membrane properties. Projecting biophysical and structural membrane and biopolymer properties to a two-dimensional plane can be beneficial to quantify molecular signatures in a reduced dimensional space to identify relevant interactions at the interface of interest, i.e. the membrane surface or biopolymer-surface interface. Here, we present a toolbox designed to project membrane and biopolymer properties to a two-dimensional plane to characterize patterns of interaction and spatial correlations between lipid-lipid and lipid-biopolymer interfaces. The toolbox contains two hubs implemented using MDAKits architecture, one for membranes and one for biopolymers, that can be used independently or together. Three case studies demonstrate the versatility of the toolbox with detailed tutorials in GitHub. The toolbox and tutorials will be periodically updated with other functionalities and resolutions to expand our understanding of the structure-function relationship of biomolecules in two-dimensions.

**SIGNIFICANCE:** 2Danalysis is a significant contribution for the systematic analysis of membrane and biopolymers simulations at planar interfaces. Developed under the open-source MDAKit framework, it enables efficient analysis of trajectories and provides versatile tools for projecting biophysical properties onto a two-dimensional plane. These capabilities enhance the visualization of spatial and temporal changes in membrane characteristics, such as order parameters, and support the study of biopolymer-surface interactions by analyzing adsorption and confinement mechanisms at 2D interfaces. The users can customize the toolbox to study complex 2D phenomena from molecular simulation.

## INTRODUCTION

Molecular dynamics (MD) simulations have significantly advanced our understanding of complex biological systems (1), including lipid membranes, proteins, nucleic acids, glycans, biocompatible polymers, and their combinations. All-atom MD is a common and powerful approach to studying such systems, propagating the motion of atoms according to Newton’s second law with empirical force fields to model atomic interactions. Simulation engines such as GROMACS (2, 3), AMBER (4, 5), CHARMM (6, 7), NAMD(8, 9), LAMMPS (10), OpenMM (11, 12), ESPREesSO MD (13), ESPResSO++(14), HOOMD-blue (15), UAMMD (16) and the ANTON machines(17–19) enable these computations, generating extensive data and trajectory files (e.g., .xtc, .mdcrd, .dcd, .dump, .dtr) that often reach hundreds of gigabytes. While trajectory analysis can be performed using analysis modules of the same engines, built-in functions frequently lack the flexibility and advanced functionality needed for sophisticated analyses. As computational capability increases, one of the current challenges in biomolecular simulation is to automatize data parsing and analysis, particularly for systems containing lipid membranes and biopolymers interacting with surfaces during adsorption, confinement, and further interfacial phenomena.

Lipid membranes are dynamic platforms that actively participate in cellular signaling cascades, protein activation, and mechanotransduction (20, 21). As barriers for the cell, the study of membrane lipids species and their role in membrane remodeling and response to external stimuli has increased (22, 23). Lipid diversity in cell membranes prompts the formation of micro-domains or lipid rafts that differ in packing and internal order, which are challenging to study due to their transient nature (24–28). MD simulations offer a valuable tool to study such systems at high spatial and temporal resolution. Typical analysis from membrane simulation studies include area per lipid, membrane thickness, deuterium order parameters, and lipid packing defects among other structural and mechanical properties; these provide insights into membrane biophysics, time correlations, and verify agreement with experimental measurements (23, 29–34). While these properties offer qualitative insights into the structure-function relationship of the simulated systems, their application has traditionally been focused on ensemble averages for the membrane bulk (35, 36). Several analysis software published over the last decade work with specific trajectory formats or output files from a specific MD engine (37–40), which is particularly limiting for beginner users. An analysis tool to capture local variations in a two-dimensional grid-based fashion can be beneficial to understand the features and forces that govern interactions at the cell membrane and its interior. In addition, a tool that enables the user to systematically determine appropriate time frames for analysis and property correlations as well as to track their evolution during the simulation.

Polymers have been widely investigated and modeled in material science, and are currently used as models for big biological systems. The systematic analysis of their structure and related properties has been systematically tackled in three-dimensions given the vast degrees of freedom these systems posses with common flexible and semi-flexible chains of monomers. In addition, several tools are embedded in molecular simulation/analysis engines (10, 13–15, 41) to calculate polymer morphological and physical properties. However, the molecular simulation analysis of (bio)polymers interacting with surfaces during adsorption (42–47), confinement (48–50) or further interfacial phenomena (51–58) has not been yet systematically tackled. Insights into the available configurational states of biopolymer residues, in particular at a planar interface of interest, constitutes a strong tool to understand dynamic behavior during biopolymer adsorption and the collective behavior of specific regions. Moreover, to interpret interfacial phenomena that occur at very close proximity, such as hydrophilicity, which is commonly difficult to characterize in 2D from standard analysis. Throughout this manuscript, the term *‘biopolymers’* comprises: proteins, nucleic acids, glycans and bio-compatible polymers.

In this work, we introduce a python toolbox designed to project structural analysis and physical properties from simulations of lipid membranes and biopolymers into a planar interface to track local changes as molecular interactions take place and morph. Packaged as an MDAKit (59) for MDAnalysis (41, 60), the toolbox is user-friendly and easy to install, and it offers the versatility of working with multiple trajectory file formats. Projecting properties of biomolecules to the 2D plane enables a comprehensive exploration of the interactions between lipid membranes and biopolymers. 2Danalysis is a robust toolbox for researchers interested in in-depth statistical analysis of structure-function relationship, molecular signature of biological interactions, and the lifetime of local variations in biophysical properties due to interactions at biomolecular interfaces.

## TOOLBOX DESCRIPTION

2Danalysis is an open source software based on Python object-oriented programming and packaged as an MDAKit for ease of access. The toolbox was created with two main objectives *(i)* study and visualization of membrane properties as projected to the 2D membrane plane, and *(ii)* characterize the behavior of biopolymers (proteins, nucleic acids, among others) in 2D as they interact with physical or biological surfaces. The following sections present the versatility and use of the current version of this toolbox, however, the analysis modules and functions will be maintained and updated periodically to benefit the biomolecular simulations community. The latest documentation and installation instructions are available at the User Guide website: https://twodanalysis.readthedocs.io/en/latest/, and detailed tutorials are published in GitHub @pyF4all (61).

To study lipid membranes, we provide the following classes: MembProp, Cumulative2D, Voronoi2D, and PackingDefects. MembProp sets the variables needed in Cumulative2D, Voronoi2D, and PackingDefects Figure 1. MembProp defines membrane atom groups and simulation box parameters, such as lipid heads and periodicity, and infers other molecular features such as atom polarity and lipid structure. The networkx library is used to infer such features and ensure proper atom connections of all lipids in the CHARMM36 force field (C36 FF) (62)(63). The user can modify force field nomenclature if needed in the MembProp code, but the initial version of the toolbox utilizes C36 FF notation by default. Cumulative2D, Voronoi2D, and PackingDefects project properties onto the two-dimensional plane of the membrane surface as explained below.

**Figure 1.**
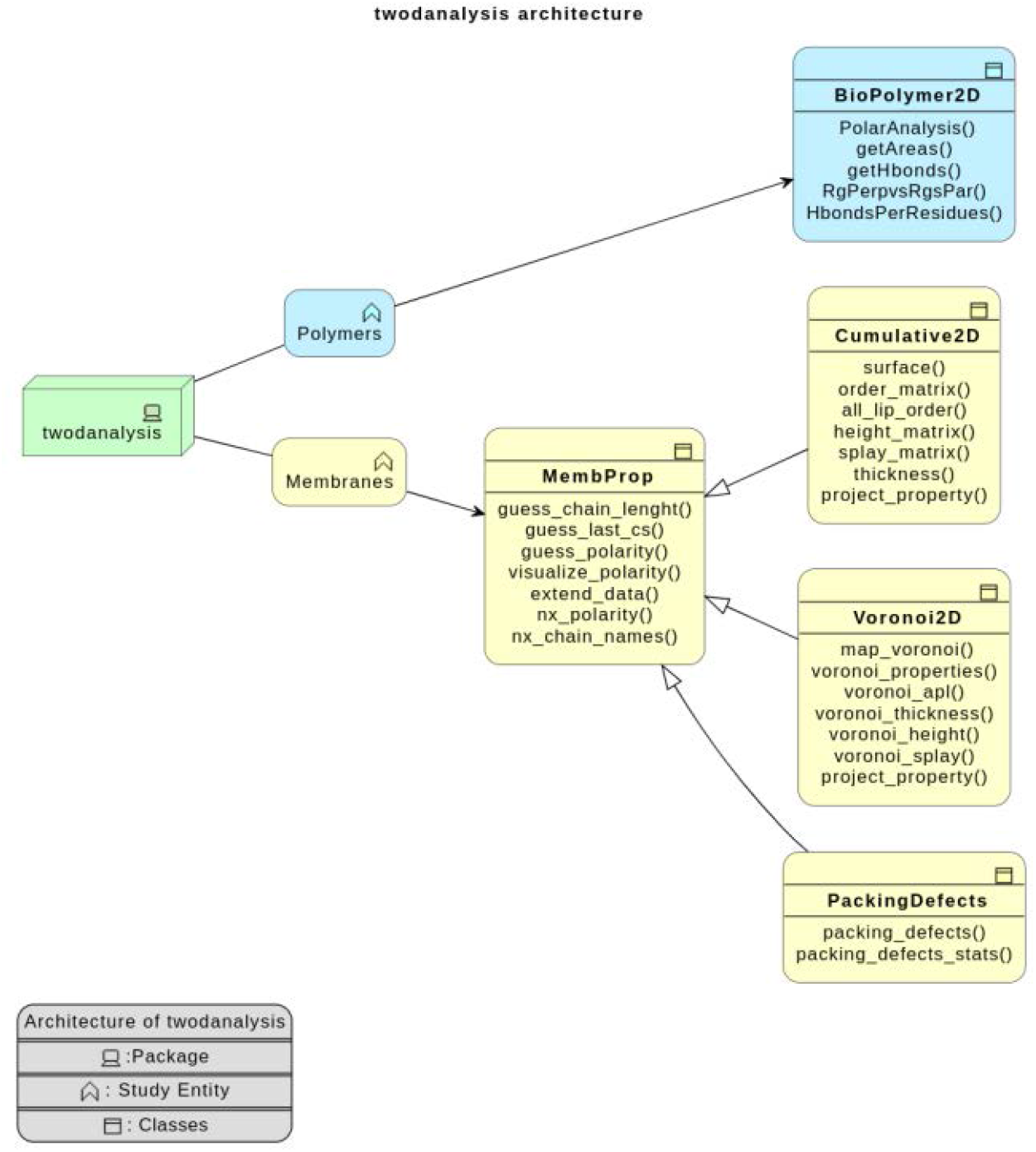
2Danalysis architecture. The blue branch focuses on analysis for biopolymers, and the yellow branch for analysis projected onto the membrane plane of lipid-only or lipid-biopolymer systems. Classes can be use independently or combined depending on the user needs. The user can use the functions contained in each class and customize the selection as needed

To characterize dynamics and interactions of biopolymers, including proteins, nucleic acids, glycans and biocompatible polymers we introduce BioPolymer2D, a class that leverages built-in functions in the MDAnalysis package to facilitate the rapid analysis of 2D interfaces between biopolymers and other surfaces. Currently, the class contains structural analysis (parallel and perpendicular radii of gyration), 2D density distribution, and polar histograms based on the center-of-mass positions of the selected biopolymer. In addition, this class performs hydrogen bonds analysis to understand the hydrophilic interactions with planar surfaces. Upon adsorption onto materials or biological surfaces, biopolymers can significantly alter their configuration, resulting in substantial changes in their electrostatic properties and free binding energies (64). Understanding these changes is essential for elucidating favorable configurations and critical residues or regions that drive or enhance biopolymer adsorption.

## METHODOLOGY

The toolbox classes are initialized by an MDAnalysis *Universe* or *AtomGroup*. In this section we describe the organization of each class and its functions for the analysis of lipid membranes and polymers.

### Study of membranes

MembProp sets the environment variables for the Cumulative2D and Voronoi2D classes, which use several functions to compute and project specific properties onto the surface plane of the membrane – albeit using distinct methodologies. Figure 2 illustrates the protocol for these projections. Cumulative2D creates a more pixelated appearance but is computationally inexpensive. Conversely, Voronoi2D produces a smoother graphical representation due to its spatially adaptive partitioning, it is computationally more expensive. Despite these differences, the overall trends captured by both methods are comparable.

**Figure 2.**
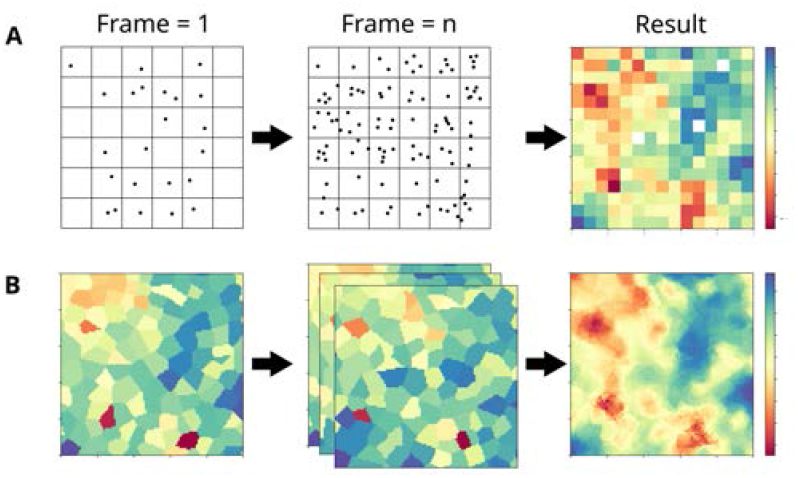
Protocols to project membrane properties to the membrane surface plane. A) Divide the space into a grid, accumulate data for *n* frames and then take grid averages. B) Generate a Voronoi diagram and map it to a 2D-grid, then compute the average over *n* frames.

For Cumulative2D, the process begins by dividing the space into a *m* × *m* grid. The positions of lipid head groups (typically phosphorus atoms) and the desired values for analysis (e.g., height) are collected over *n* frames. These values are then averaged within each grid square, thus producing a *m* × *m* matrix that contains the averaged property values. Alongside this matrix, the grid edges are recorded in the format [*x*_min_, *x*_max_, *y*_min_, *y*_max_]. Figure 2, Panel A, depicts the process.

In contrast, Voronoi2D first constructs a Voronoi diagram using the positions of the lipid head groups (typically phosphorus atoms), which is mapped onto a *m* × *m* grid. During the mapping step, the value of the desired property is assigned to the grid squares corresponding to each lipid. This mapped grid is created for each frame, as illustrated in Panel B of Figure 2, and then averaged across *n* frames. The output, similar to Cumulative2D, is a matrix *m* × *m*, along with the edges *x*_min_, *x*_max_, *y*_min_, *y*_max_ .

Either approach can be used to project the following properties to the two-dimensional membrane surface plane:

#### Membrane Thickness

can be calculated by calling the functions `thickness()` or `voronoi_thickness()` from Cumulative2D and Voronoi2D, respectively. Although both provide similar output matrices, they differ in the projection method and on the definition of membrane thickness itself. Cumulative2D computes the thickness by obtaining the mean height from the middle of the membrane for *n* frames (`Cumulative2D.height_matrix()`) for each membrane leaflet and adding them to obtain the final result. On the other hand, Voronoi2D computes the height matrices for the two layers and sums them for each frame; the final result is obtained by averaging all the matrices that computed the thickness on a per-frame basis.

#### Membrane Order Parameters

enables the projection of local order parameters onto the membrane plane, instead of the membrane-bulk average values. This structural property is lipid-dependent due to variations in the fatty acid tail length and chemical structure. For each lipid, the deuterium order parameter (*S*_*CD*_) value is computed using

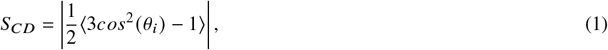

where *θ*_*i*_ is the angle between the C-H vector of the *i* ™ th carbon and the membrane normal (z-axis). The calculation considers both tails (*sn*1 and *sn*2) of the lipid, yielding *m* + *n* values, where *m* and *n* represent the number of carbons in *sn*1 and *sn*2 respectively. Using the standard protocol in Cumulative2D, these values can be projected onto the membrane surface plane for spatial analysis. This process involves creating *m* + *n* matrices –one for each carbon in the lipid– that contains the averages for each grid square in the two dimensional plane. The first *m* matrices are averaged (corresponding to *sn*1); then, the *n* following matrices (corresponding to *sn*2); subsequently, the results for each lipid tail are averaged to obtain a single matrix for *S*_*CD*_ per lipid (`Cumulative2D.order_matrix()`). Finally, the results are averaged over all lipids for a complete 2D representation on the membrane plane (`Cumulative2D.all_lip_order()`). Currently, two dimensional projection of *S*_*CD*_ is not available for Voronoi2D.

#### Membrane properties

current analysis functions in the classes above include *Height* of lipid headgroups measured from the middle of the membrane, *Splay angle* between lipid tails, and *Area per lipid* that the user can directly plot onto the membrane surface plane. In addition, the user can compute custom properties and use the Cumulative2D and Voronoi2D classes to plot them in 2D. To this end, the user must first define a function that takes as input an atom selection from MDAnalysis, `mda.AtomGroup`, and returns two numpy arrays. The first array must store the lipid residue indexes (`mda.AtomGroup.residues.resids`), while the second array stores the property value of interest for the respective lipid molecule as listed by index in the first array. The userdefined function serves as an argument to `Cumulative2D.project_property()` or `Voronoi2D.project_property()` which will apply the *Cumulative* or *Voronoi* approaches to render the projections, respectively.

#### Packing defects

based on PackMem (65), which maps hydrophobic and hydrophilic atoms onto a 2D grid according to their radii, our implementation provides enhanced code performance by computing small matrices for unique atomic radii and storing them in a dictionary at initialization to eliminate redundant comparisons for atoms that share the same radius. PackMem computes the area of packing defects by counting the grid squares with values less than or equal to 1, values that demark hydrophobic and hydrophilic regions, respectively. Their recommended grid size of 1 lies in a regions where the packing defect area is still evolving, as shown in Figure S1. Since the computed area is not at equilibrium, the computed value may not be converged and result in larger uncertainties. Finer gid sizes improve precision, but they substantially increase computational costs in PackMem. 2Danalysis mitigates this computational overhead and enables the use of smaller grid sizes (0.5 or finer) with much smaller impact on performance. Additionally, we address periodic boundary conditions by replicating 10% of the data at each edge of the grid, ensuring continuity across the simulation box boundaries. PackingDefects.packing_defects() computes packing defects for a single frame, and returns a 2D matrix with the packing defects and a dictionary containing information about the defects such as size, defect area, and matrix edges among others.

### Study of Biopolymers

BioPolymer2D contains four functions to characterize the adsorption and confinement mechanisms of biopolymers onto surfaces: *(i)* parallel (*R*_*g*∥_) and perpendicular (*R*_*g*⊥_) radii of gyration; *(ii)* polar histogram; *(iii)* 2D-projected density; and *(iv)* H-bonds per residue/nucleotide/monomer. These routines aim to facilitate the study of complex interfacial phenomena that modulates and stabilizes biopolymer adsorption.

These functions consider absorbed states of the biopolymer during the simulation trajectory, as determined by a distance threshold from the surface of interest (`zlim`) and the number of simulation frames that comply with this condition (`Nframes`) . Note that the distance threshold will also define the total Nframes included for the analyzed trajectory. All the routines in BioPolymer2D use the term *residue* to denote protein amino acids, nucleic acid nucleotides, and individual monomers as it pertains to the system under study.

#### Polar histograms

computes the distance of individual residues to the center of mass of a reference selection of atoms. This type of analysis gives detailed information of the residues positional fluctuations over the simulation trajectory. These histograms can be interpreted as the probability distribution of each residue; therefore, the width of the histogram is indicative of the looseness of a *residue* with respect to the planar surface. This type of analysis has been previously introduced in ref. (42)

#### 2D-projected density

computes a two-dimensional density distribution of the positions of a group of residues using the KDE method in the Seaborn Python package (66, 67). The 2D-projected density identifies the residues that exhibit the highest interaction at the planar interface of interest according to a user-set criteria. Similar to the *Polar histograms*, the area of the contour plots informs on residue looseness or frequency of contact. The user can set density thresholds to examine specific interactions and compute the area spanned by the residue(s) during the analysis timeframe. On the one hand, the area of the lower contour levels, samples all the available space states in the both surface axes (e.g. X and Y). This can be interpreted as an estimation of adsorption looseness. On the other hand, by computing the area of a higher contour level, we are able to sample the peaks of 2D density distribution, which is related to the strong and specific interactions with the surfaces.

#### Parallel and perpendicular radii of gyration

computes the standard 3D (*R*_*g*_), parallel (*R*_*g*∥_) and perpendicular (*R*_*g*⊥_) radii of gyration.

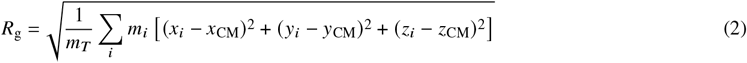

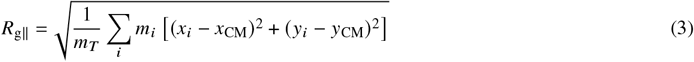

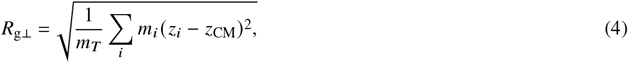

where **R**_CM_ = *x*_CM_, *y*_CM_, *z*_CM_ is the position of the center of mass, *m*_*i*_ the mass of each residue and *m*_*T*_ the total mass of the residues.

The parallel and perpendicular radii of gyration give structural information of the adsorption phenomenology. Namely, *R*_*g*∥_ gives information on how the biopolymer is deformed by the sides (parallel to the surface), and *R*_*g*⊥_ offers a measure on how the biopolymer is stretched or flattened perpendicular to the surface.

On top of calculating *R*_*g*∥_ and *R*_*g*⊥_, BioPolymer2D generate figures showing *R*_*g*⊥_ vs *R*_*g*∥_ values, and indicating 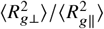 ratio, which is a well studied criteria for tackling polymer adsorption (68) and has been recently adapted to RNA (43), proteins (42) and a combination of proteins and glycans, as shown in Figure 6A.

## CASE STUDIES

The examples below illustrate prominent features of the 2Danalysis toolbox. The first case study focuses on membrane properties for a complex lipid mixture whit a nucleic acid fragment that interacts with the surface. The second case examines a complex membrane model in contact with a peripheral membrane protein. To conclude, the third case explores a protein-glycan pair adsorbed onto a generic surface. Specific details of the simulation settings for the respective trajectories can be found in the supplementary material. While these showcase many capabilities, the toolbox offers more features and versatility that are detailed in the tutorials available in GitHub @pyF4all(61).

### Case study 1: lipid-lipid interactions from a membrane-RNA simulation

Figure 3 presents analysis performed using the MembProp class on a membrane-RNA system, with primary focus on membrane biophysical properties projected to the membrane interface plane. The membrane model containing DSPC, POPE, DODMA, CHL lipids (45:20:15:20 mol %) was build using CHARMM-GUI Membrane builder (69–71) (Figure 3F). The system included an RNA (72) fragment in the water and neutralizing ions, and was simulated for 500 ns with the C36m FF (62, 63, 73) on GROMACS using the NPT ensemble at standard conditions for fluid membranes. Figure 3 A-C show images rendered using the `Voronoi2D` (left) and `Cumulative2D` (right) classes to contrast their resolution for the analysis of 50 frames of the simulation trajectory stored at 0.5 ns intervals. The increase in resolution for the left panels is a direct consequence of mapping the Voronoi diagram onto a 2D grid, which allows the use of more bins (`nbins=180` for this example). For the plots generated with `Cumulative2D` in Figure 3, we used `nbins=50`. On the other hand, using higher number of bins with `Cumulative2D` results in grid spaces without data to average, preventing the rendering of the image. Despite this difference in resolution, both classes show similar trends.

**Figure 3.**
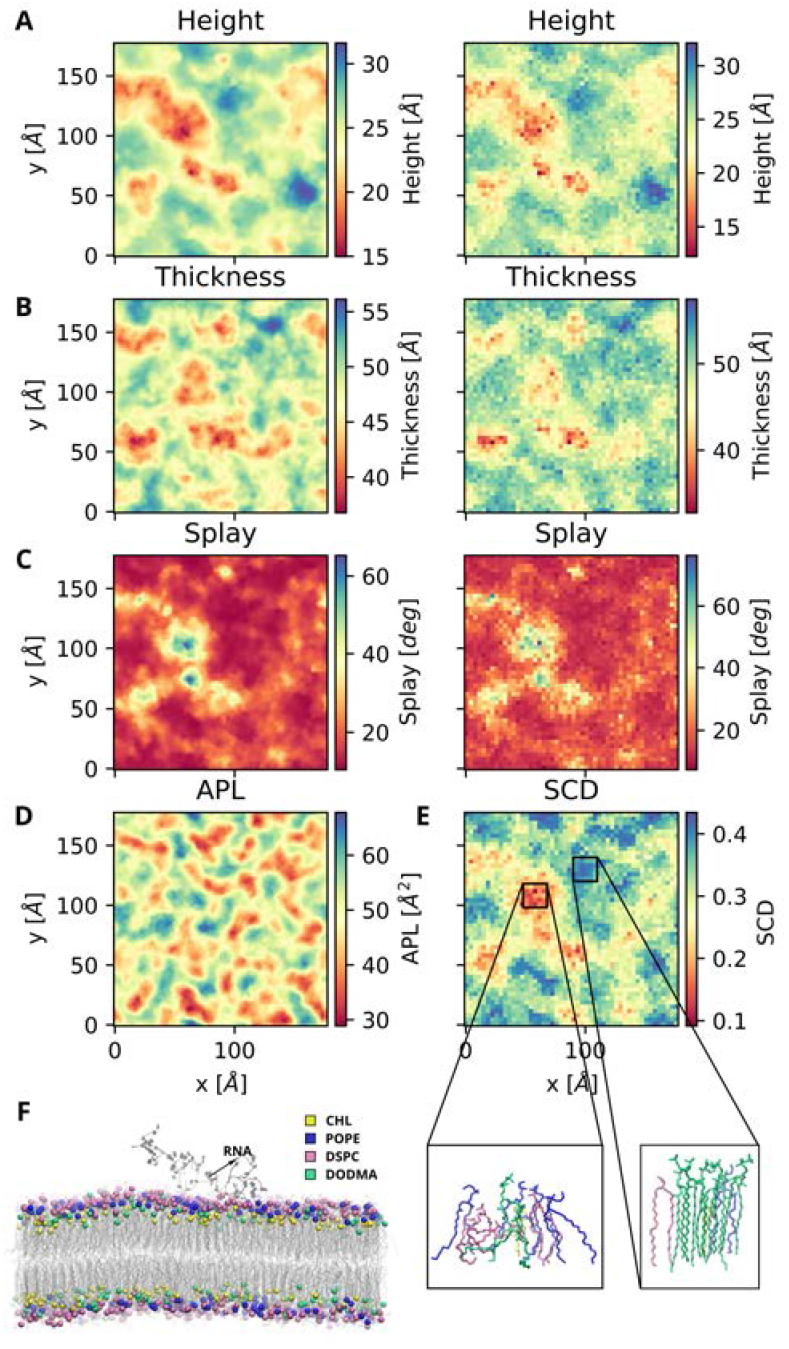
(A-C) Plots obtained using Voronoi2D (left) and Cumulative2D (right) for a DSPC:POPE:DODMA:CHL (45:20:15:20 mol %) lipid mixture for 25ns between the 30-55ns of simulated trajectory. (A) Membrane height defined using the average z-position of *P* lipids in the top leaflet. (B) Membrane thickness. (C) Splay angle for lipids in the top leaflet. (D) Area per lipid for the top leaflet. (E) Deuterium order parameters for lipids in the top leaflet showing snapshots of local lipids in the regions with lower and higher order, respectively. (F) Lateral view of the lipid-RNA system.

The main advantage of studying membrane properties with a two-dimensional approach is the ease of identifying regions that may be related to biological or physical phenomena. Splay angle, membrane thickness, APL, and SCD are related to line tension (74), bending rigidity (75), lipid-raft formation (74), ion permeation, membrane disruption(76, 77), and liquid-ordered and liquid-disordered phases (78, 79) among others. Figure 3A-E show regions with different property values; identifying these regions in the 2D plane allow us to examine those areas more closely to determine the molecular interactions as well as chemical environment that give rise to the change in property value. For example, Figure 3E shows membrane regions with lower (red) and higher (blue) projected *S*_*CD*_ values. Snapshots of the lipids at the corresponding coordinates rendered on VMD (80, 81) show lower order regions correspond to entangled lipid tails that exhibit larger splay angles, even for lipids like DSCP and POPE that are tend to exhibit more ordered behavior. On the other hand, regions with higher *S*_*CD*_ host lipid tails vertically aligned to each other and narrower splay angle, even for DODMA lipids that tend to intertwined with their neighbors.

Some membrane properties can exhibit correlation, like the surface topology determined from the analysis of height per leaflet and the membrane internal structure using the lipid order parameters (*S*_*CD*_). 2Danalysis allows the quantitative examination of such relationship by quantifying correlations between specific features. Figure 4 C shows a nearly linear relationship between *S*_*CD*_ and membrane thickness, a well-known relationship that was first introduced by Seelig and Seelig (82, 83). The original work considered a pure DPPC bilayer, here we show this trend is also true for a complex lipid mixture. It is important to note that the correlation degree and direction may vary for other lipid compositions; data spread, simulation temperature and pressure, and ring moieties in lipid molecules are some of the factors that may affect this correspondence. 2Danalysis outputs facilitate the pipeline to quantitative analysis of 2D-projected data on a per-grid basis.

**Figure 4.**
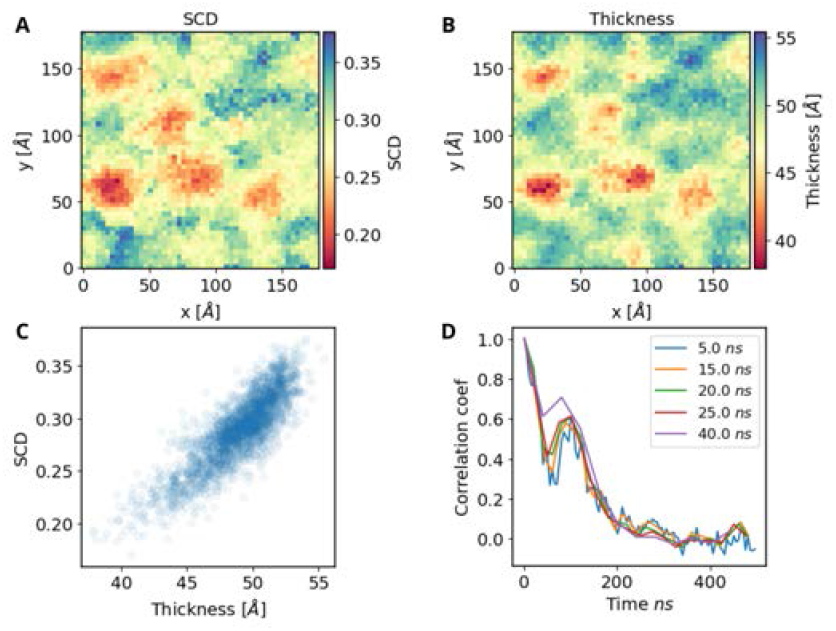
Spatial similarity between (A) *S*_*CD*_ values, and (B) membrane thickness. (C) Scatter plot to depict the correlation of *S*_*CD*_ vs thickness per grid square. (D) Correlation coefficient of the matrices with varying time window size for the same trajectory

As mentioned before, local membrane properties change over time as lipids and other biomolecules interact and diffuse laterally. Understanding how fast this properties change is critical to characterize membrane dynamics. A quantitative analysis of the rate of change of a given property can also aide in the systematic selection of time-windows for specific analyses. Figure 4D shows the correlation index of *membrane thickness* over time using increasing time-windows. A convolution with a normalized 3 × 3 kernel is applied to individual matrices to asses their similarity over time computing the Pearson correlation for the

normalized matrices. The correlation coefficient for membrane thickness follows the trend of an exponential decay, reaching its half-life at approximately 40 *ns*. Therefore, for the purposes of comparing different portions the simulation trajectory, the optimal analysis time-window should be 40*ns*. Note this characteristic time may vary among membrane properties, and should be considered carefully depending on the metric(s) that best describe the process of interest.

### Case study 2: lipid membrane-protein simulations

The previous case study discussed examples considering only lipid-lipid interactions. 2Danalysis is an ideal tool to study lipid interactions with other biomolecules since it easily identifies local and global changes in membrane properties both quantitatively and quantitatively. This case study leverages 2Danalysis to expand on a study of the Mixed Lineage Kinase Like (MLKL) murine protein with a model for the plasma membrane previously published in (32). The membrane model is composed of a mixture of five lipids (DOPC, DOPE, CHL, PIP, PIP_2_) with the protein positioned 1-2nm above the bilayer in different orientations. (Figure 5) expands on the published analysis using the third replica of the original study in which the protein interacts with the membrane through its four-helical bundle (4HB) and brace regions (see snapshot inFigure 5B).

**Figure 5.**
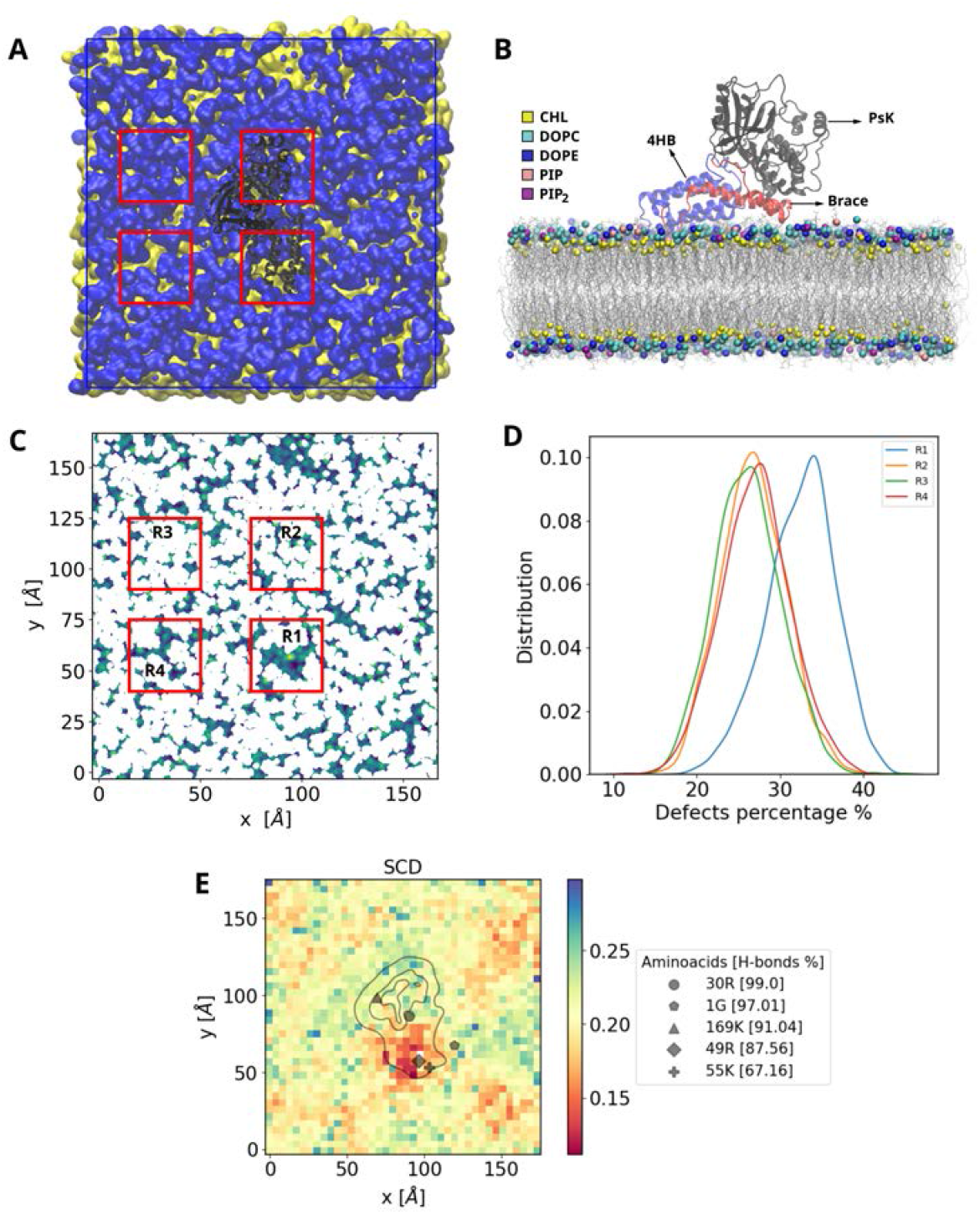
MLKL-membrane system. (A) Top view of the system with the protein in black, hydrophobic atoms in yellow and hydrophilic atoms in blue (B) Lateral view of the system (C) Packing defects of the binding leaflet obtained with `PackingDefects.packing_defects()` (D) Probability distribution of the percentage of defects in different regions of the binding leaflet

In the original study, packing defects were computed using the protocol described by Wildermuth et al. (84), which is computationally intensive as it requires rendering and processing multiple images of the membrane surface at small time intervals. The yellow regions in Figure 5A represent the packing defects identified using that method, while Figure 5C shows the same analysis performed using `PackingDefects.packing_defects()`; both approaches render close results as seen in the boxed portions.

The `PackingDefects` class allows for detailed comparisons among specific regions of packing defects. For example, Figure 5C shows four regions in the membrane surface. R1 corresponds to the area where the 4HB and brace domains of the protein interact with the membrane, while the other regions were selected randomly for comparison. The packing defect percentage in R1 is notably higher compared to the randomly chosen regions (see the distributions in Figure 5D). Visual examination of the trajectory reveals the increase in packing defects is due to the interaction and partial insertion of the 4HB into the bilayer. This is in alignment with the original findings in (32), but allows a quantitative comparison of the progressive increase of packing defect area at the protein binding site.

Finally, Figure 5E highlights the power of combining two classes, `Cumulative2D` and `Biopolymer2D`, to study protein-lipid interactions. The membrane background was rendered using `Cumulative2D.all_lip_order()` to compute the *S*_*CD*_ of the binding leaflet. Overlaid on the membrane map is a contour representation of the protein and key residues forming hydrogen bonds with membrane lipids. The greater contributors to hydrogen bonding are protein residues in the 4HB interacting with lipids in a region with low order (shown in red). 2Danalysis allows a holistic perspective of the system, examining local membrane properties and protein-membrane interactions in tandem. Results of the method are returned in the form of 2D matrices, which allows for further quantitative analysis using different time windows to track changes at the protein-lipid interface.

### Case study 3: Adsorption of a SARS-CoV2 RBD in closed conformation onto a planar surface

Using the BioPolymer2D class, we systematically characterize the interaction of the RBD of the SARS-CoV2 omicron variant and a glycan on a hydrophilic surface; this class is specially useful to determine flexible regions in the biopolymers. Glycans are known to be more flexible than proteins, because they have a single covalent bond to proteins and do not acquire any secondary structure. In Figure 6, we analyze the looseness and structural characteristics of this protein-glycan complex at the interface of a hydrophilic surface.

**Figure 6.**
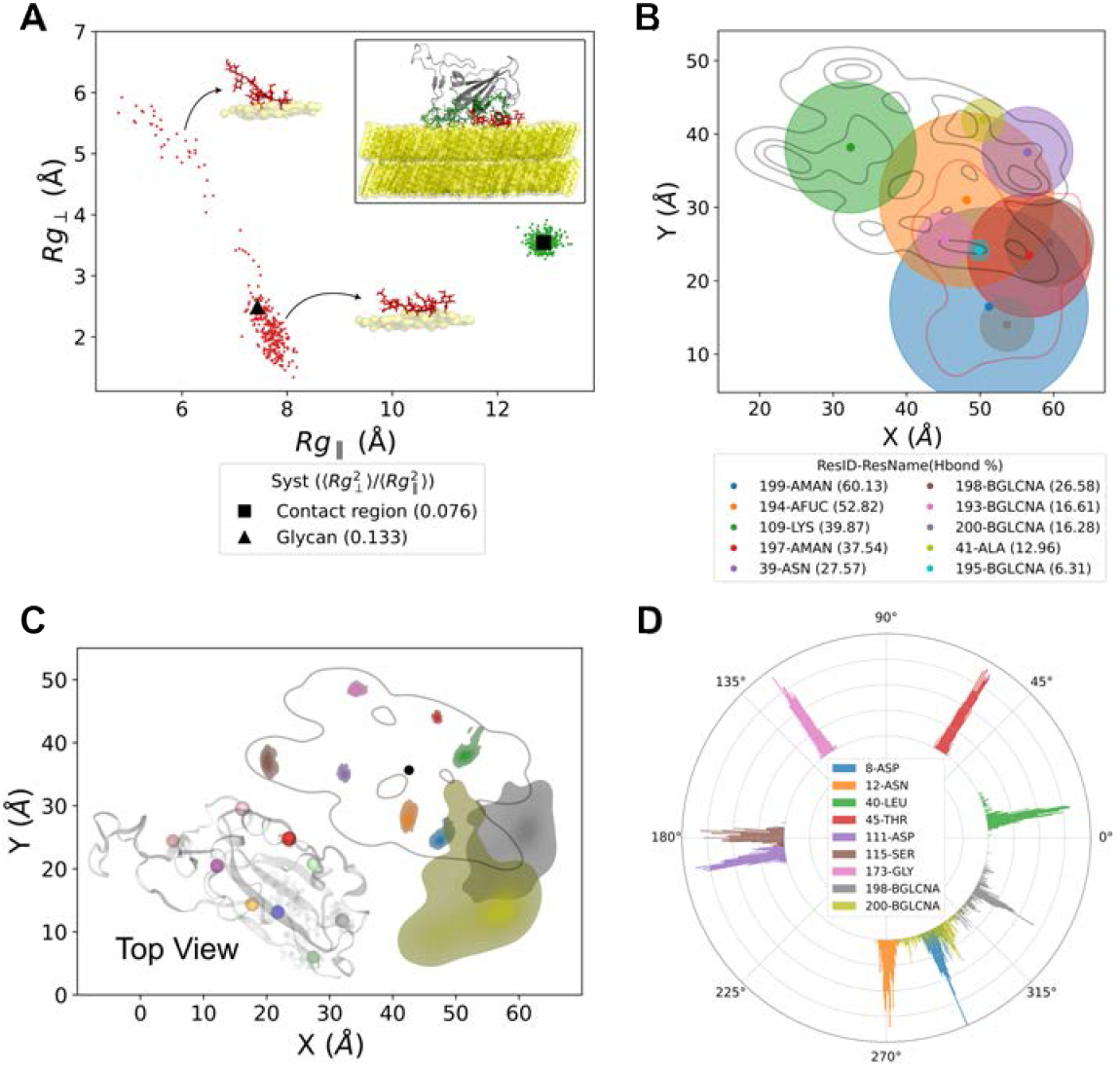
Protein-glycan complex on a hydrophilic surface.(A) Parallel and perpendicular radii of gyration of the glycan (in red) and adsorbed protein residues (in green) during the full trajectory. Snapshots of the glycan in different configurations are shown as reference. The mean *R*_*g*∥_ and *R*_*g*⊥_ values of the asborbed biopolymers are marked by a triangle and square, respectively. (B) Highest occurrence of hydrogen bonds per biopolymer residues during the trajectory (in %) shown on a top view of KDE contour of the protein-glycan complex and the representative areas for the ocurrence of H-bonds per residue in the top ten. (C) 2D density distributions (protein contour and snapshot shown as reference) and areas of the residues with highest contacts with the surface; and a (D) Polar histogram showing the looseness between the protein-glycan complex and the surface, references on the complexes center-of-mass; the color legend in D applies to the bottom panels

The parallel and perpendicular radii of gyration (see Equation 3 and Equation 4) of the protein and the glycan were computed separately using the `getRgs2D` function. This function can also compute the 3D radius of gyration of an atom selection using (Equation Equation 2), to plot the time series of the three Rgs set the input parameter `plot=True` (see Figure S2). The ratio between *R*_*g*∥_ and *R*_*g*⊥_ can also be computed for further analysis of the lifetime of specific configurations, and displayed on the plot setting the input `RgPerpvsRgsPar`. Note the contrast between the data for the glycan (red) and the protein residues (green) interacting with the hydrophilic surface in Figure 6A for both *R*_*g*∥_ and *R*_*g*⊥_. The scattered points show an enhanced looseness for the glycan while adsorbing onto the surface. Interestingly, the 3D radius of gyration alone cannot identify such dynamic spatial behavior (see Figure S2). As for the contact region of the protein, we can evidence that all data points in green are surrounding the mean value (see Figure 6A). This could indicate that there is not much looseness during protein adsorption despite having a large mean *R*_*g*∥_ value. In fact, the protein retains its secondary structure intact during adsorption, as shown in Figure S3, where the variation of different secondary structure categories is negligible (less than 3% at it peaks) during the simulation trajectory.

The H-bonds of protein and glycan residues at interface with the hydrophilic surface can be calculated using the BioPolymer2D function `getHbonds`, and show the H-bonds with `plotHbondsPerResidues` method. Figure 6B shows an example of the plots this function can generate, where the percentage of H-bond formations along a trajectory by increasing proportionally the area of a circumference. The legend of Figure 6B, shows the top ten residues or glycans forming H-bonds during a trajectory. In this figure, we can evidence that the Glycans play a determinant role in the adsorption by forming H-bonds with the surface over several top values in the ranking 60.13%, 52.82% and 37.54% of the trajectory. Moreover, all glycan residues are part of the top 10 residues with the most H-bonds.

To further characterize the looseness of the biopolymers during adsorption, use the Polar histograms (`PolarAnalysis`) and 2D-project density (`KDEAnalysisSelection`) functions to analyze the areas of interaction between protein and glycan residues that exhibit most contact with the hydrophilic surface. In Figure 6 C-D, the glycan residues exhibit a large increase of contact area as shown by the respective outer contour levels. Moreover, using the `getAreas` function, which uses the Simpson integration algorithm, we can estimate the surface area of the desired contour levels. Table 1, lists these areas for the colored residues in Figure 6C, evidencing that contact areas of glycan residues are approximately one order of magnitude higher than those of the adsorbed protein residues. Likewise, Figure 6D shows the enhance looseness of the Glycans-surface with respect to the protein-surface interactions, observed by the wider distribution of residue fluctuations with respect to the center-of-mass of the combined selection of atoms for the adsorbed protein-glycan complex.

**Table 1:** Outer level areas per biopolymer residue computed from the contour plots in Figure 6C.

| ResIDs | Resnames | Area (Å<sup>2</sup>) |
| --- | --- | --- |
| 8 | ASP | 3.70 |
| 12 | ASN | 3.55 |
| 40 | LEU | 2.80 |
| 45 | THR | 0.66 |
| 111 | ASP | 1.83 |
| 115 | SER | 4.71 |
| 173 | GLY | 2.34 |
| 198 | BGLCNA* | 56.05 |
| 200 | BGLCNA* | 68.94 |

\* Glycan residues.

## DISCUSSION

2Danalysis is a toolbox that offers versatility to new and experienced users of MD simulations to study biomolecular interactions. It is a Python object-oriented open source software packaged as an MDAKit in MDAnalysis for ease of access and compatibility with all the most common trajectory file formats from molecular simulations. The main goal of this toolbox is to reduce the dimensionality of large simulation data sets to a two-dimensional plane of interest. For example, the lipid membrane surface, the binding interface between two biomolecules, and the adsorption plane between a (bio)molecule and a surface. This toolbox is versatile and powerful to examine local changes that result from the interactions of (bio)molecules, and how these change the local molecular landscape in a planar interface. 2Danalysis is organized into classes to facilitate the analysis of membrane systems (MembProp, Cumulative2D, Voronoi2D, and PackingDefects) and biopolymers (BioPolymer2D), which can be used separately or in tandem.

Through three case studies, we demonstrated the toolbox’s utility to explore properties in 2D for lipid-lipid, lipid-biopolymer, and biopolymer-surface systems. The toolbox facilitates the computation of typical biophysical membrane properties as well as versatility for user-defined functions that leverage atom selection syntax from MDAnalysis and the outputs from class functions of 2Danalysis. The user can compute correlation indices to evaluate the similarity across different time windows and examine phenomena that may occur within specific temporal ranges, and utilize the suggested statistics-based approach to quantify spatial correlations between properties of interest.

The class to compute lipid packing defects significantly improves the efficiency of previous methods (65, 84) to compute this metric, and allows the user to define a finer grid to reduce statistical error. The user can focus on specific membrane regions to study local effects of biopolymer adsorption on the surface area of packing defects. For instance, the outputs of `PackingDefects.packing_defects()` are a matrix and a dictionary with information on individual defects, which can be used to estimate depth of individual defects using the corresponding value for the density (number) of hydrophobic atoms in that region.

Outputs from the MembProp class serve as input variables for the functions in the `Voronoi2D` and `Cumulative2D` classes, see Table S1. These can be readily applied for the analysis of membranes with any lipid composition. The user can systematically select and customize the appropriate time frame for analysis at different trajectory points for comparison, and update reference atoms for the selected analysis as needed. In case study 2 we introduce one approach to study spatial correlations on a per-grid-square basis. Other ways to explore spatial correlation metrics, such as Moran’s I index, are discussed in (85–87).

BioPolymer2D functions extend on established 2D structural analysis of (bio)polymers (42, 43, 48, 88) and feature new analysis to understand individual local interactions at the interface of biopolymers-surfaces. In case study 3, we tackle a biological system of a protein with its corresponding glycan, where the flexible character of the glycan and semi-flexible one of the protein during adsorption can be identified by the areas coming from the 2D-density distributions and histogram of the polar plots. Moreover the H-Bonds formed during adsorption provide an overview of the per-residue hydrophilic character of the short-range interactions. On top of this, based on the characteristics of the 2D-density analysis, this toolbox is able to group several residues/atoms and perform local structural analysis of *R*_*g*_ radii to elucidate *e*.*g*. domains with loosen or tight adsorption patterns (42). Such a combined analysis can also be exploited to characterize the effect of mutations (42) or that of contaminated surfaces (58) at interfacial domains during biopolymer adsorption or confinement. The combination of *R*_*g*_ radii, polar analysis and 2D-density distributions analyses provides a powerful tool for distinguishing the adsorption patterns (trains, loops and tails) (89) of any biopolymer at planar interfaces.

The functions in 2Danalysis offer efficient, versatile, and systematic analysis, expanding on existing work(37–40), and focusing on interactions that occur in two-dimensional planes. The potential of a systematic and automatized analysis of all-atom interfaces stored in consolidated data projects (90) improves statistics and hence better models of interfacial biophysics. For instance, the versatile character of this tool enables its integration to other molecular resolutions (spatial scales), which can include available coarse-grained biomolecular force fields (91–95). This tool offers a pipeline for the systematic comparison between coarse-grained molecular models and their original all-atom representation, *e*.*g*. protein residues modeled as individual beads. Finally, 2Danalysis returns outputs as 2D matrices that can be easily fed to Machine/Deep Learning models to uncover important regions in the membrane or at the surfaces.

## CONCLUSION

The development of this toolbox represents a significant contribution in the study of membranes and biopolymers interacting with materials or biological interfaces. Specifically, our tools provide an efficient way to identify and analyze quantitatively molecular interactions in two-dimensional planes, enabling the analysis of statistical correlations between structural and dynamic properties of lipid membranes and biopolymers. Released as an open-source project under the MDAKit framework, it invites researchers to explore, modify, and extend its functionality, supported by comprehensive documentation. By enabling robust and streamlined analyses, the toolbox facilitates the study of complex systems and offers a practical solution for building pipelines to efficiently analyze multiple replicates quantitatively. Users can access the latest documentation and installation instructions at the User Guide website: https://twodanalysis.readthedocs.io/en/latest/, and follow detailed tutorials published in GitHub @pyF4all (61). These sites will be maintained to incorporate current examples of the methods and classes of the latest version of the toolbox.

## AUTHOR CONTRIBUTIONS

**RR:** contributed to the development of analytic tools; code testing; integration to MDAKits; literature research. **AB:** contributed to the development of analytic tools; code testing. **RP:** graduate training and supervision. **HVG:** conceptualization; designed research; graduate training and supervision. **VM:** conceptualization; designed research; graduate training and supervision. **All authors** contributed to the writing of the manuscript

## ACKNOWLEDGMENTS

**R.R**. and **V.M** acknowledge the computational resources available through the University at Buffalo Center for Computational Research (96) and the Anton2 machine, provided by Pittsburgh Supercomputing Center (PSC) through Grant R01GM116961 from the National Institutes of Health, specific award MCB200093P. The Anton2 machine as PSC was generously made available by DE Shaw Research (17). **H.V.G**. acknowledges financial support from the Ramon y Cajal grant No. RYC2022-038082-I and Spanish Ministry of Science and Innovation, through project PIDPID2023-150536NA-I00, and the “Severo Ochoa” Grant No. CEX2023-001263-S for Centers of Excellence; and thanks the Red Española de Supercomputación (RES) for the computing time and technical support at the Finisterrae III supercomputer project FI-2024-3-0033. **R.P**. acknowledges support from the Spanish Ministry of Science and Innovation, through project PID2023–149150OB-I00, and the “María de Maeztu” Programme for Units of Excellence in R&D (CEX2023–001316-M). **A.B**. acknowledges the financial support of Comunidad de Madrid through the predoctoral research contract PIPF-2023/TEC-29920. **The authors** thank Willy Menacho, Jinhui Li, and Oluwatoyin Campbell for testing the code before its release and feedback during the writing of this manuscript.

## SUPPLEMENTARY MATERIAL

An online supplement to this article can be found by visiting BJ Online at http://www.biophysj.org.

