## Supplementary Material for "2Danalysis: A toolbox for analysis of lipid membranes and biopolymers in two-dimensional space"

### Simulation details for case studies 1-3

Examples shown in figures 3-6 are detailed in the full tutorial for the MembProp and BioPolymer2D classes [1]. Visual Molecular Dynamics (VMD) package [2] was used to visualize the trajectories and render all the snapshots for this work. Below we provide a summary of the simulation settings and building steps for each system discussed in the main manuscript. All the simulated systems were run using periodic boundary conditions.

***Case study 1:*** a symmetric lipid bilayer containing DSPC, POPE, DODMA, CHL lipids (45:20:15:20 mol %) was build using CHARMM-GUI Membrane builder [3–6], and relaxed following the CHARMM-GUI 6-step relaxation protocol. A 100ns equilibration trajectory was completed before merging the bilayer coordinates with equilibrated coordinates of a 43-nucleotide RNA fragment. This fragment was solvated using CHARMM-GUI Solution Builder [7] and equilibrated for 50ns before placing it near the membrane surface to sample unbiased adsorption dynamics. A solution of 0.15 mol/L KCl neutralizing ions was used to neutralize the final membrane-RNA simulation box of 20x20 nm, followed by an energy minimization using the steepest descent algorithm and a 500ns simulation trajectory without any molecular restrains.

All-atom molecular dynamics were performed using the Charmm36m force field [8] and a three-site water model used for solvent [9] in the NPT ensemble at 303.5K and 1 bar using the Nose-Hoover thermostat [10, 11] and Parinello-Rahman barostat [12, 13] in GROMACS 2021 software package [14]. The LINCS algorithm was used to constrain hydrogen bonds during the simulation [15]. Electrostatics were evaluated using the Particle Mesh Ewald method [16] with a 1.2 nm cutoff, and long-range interactions modeled using a van der Waals potential with a force-switching function between 1-1.2nm [17].

***Case study 2:*** the analysis presented in this manuscript is an extension of the work published in [18], refer to the main manuscript for full details of the simulation methods. In brief, coordinates of the MLKL protein (PDBID: 4BTF) and a model bilayer for the plasma membrane of eukaryotes were used to characterize the driving forces and molecular signature of protein-lipid interactions during late-stage necroptosis. The membrane model contained a mixture of DOPC:Chol:DOPE:POPI-1,4:POPI-2,5 lipids (40:32:20:4:4 mol%) and was built using CHARMM-GUI *Membrane Builder* [3–6]. The protein was solvated in water using the *Solution Builder* [7] . Equilibrated coordinates of membrane and protein were merged for a final systems with 600 lipids per leaflet and the protein in

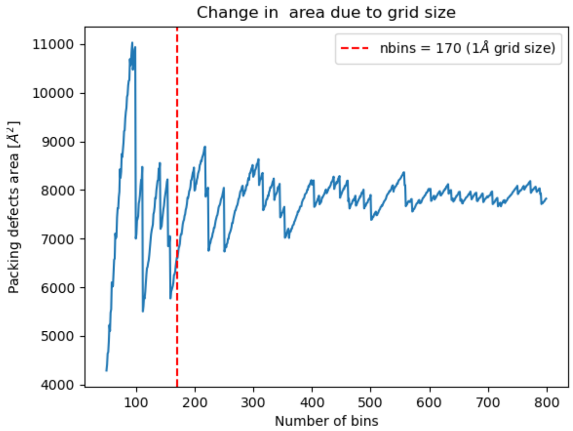


Figure S1: Packing defects area for the same simulation frame of the MLKL-membrane trajectory using different values of *nbins*. The red line denotes the 1Å grid size, which is too small to compute average packing defects since the region still retains large fluctuations in the defect area. A better value for this system to prevent loss of efficiency in the code is lower than 0.5Å

different orientations hovering 1-2nm above the bilayer in a simulation box of 17.3×17.3×18.4 nm. The full study examined four replicas, each run for 100ns in Gromacs [14] prior to transferring them to the Anton2 machine [19] for 2,000 ns trajectories per replica. All systems used the Charmm36m force field [8] with typical settings for fluid bilayers in Gromacs, and optimized parameters or the integration algorithms on the Anton2 machine that are set by its internal guesser, the *Multigrator* integrator [20].

***Case study 3:*** all-atom simulations were carried in Gromacs 2023 [21] for a system containing the SARS-CoV-2 RBD protein conformation from our previous work [22], neutralizing ions, and a polarizable modeled surface with the CHARMM36 [23, 24] forcefield, and TIP3P [25] model for water. CHARMM-GUI was used to attach the glycan to the RBD. The hydrophilic surface (PBL) was built from a small patch of decanol (DOL) with positional restraints to maintain the bilayer shape and avoid interference of the mechanical properties of the *bilayer*surface, as discussed in previous works [26–28]. The surface model is not flexible and does not allow molecular defects. The PBL surface was replicated using the built-in *gmx editconf* Gromacs program and solvated with water for a final rectangular box of 8.6×7.6×12 nm. The polarity of the OH- groups of the DOL chains was set to 1 for the hydrophilic surface [26]. The coordinates for the SARS-CoV-2 Omicron variant RBD-glycan complex were added to the box.

Energy minimization was done using the steepest descent algorithm for 50000 steps with an integration time step of 0.01 ps. System equilibration was done using the NVT and NPT ensembles for 100ps, respectively. Positional restrains of 250 kJ $mol^{-1}$ $nm^{-2}$ were applied to residues in close contacts between the RBD and S1 regions of the protein along the $x$ and $y$ axes, while retaining the flexibility of the remaining protein residues and fluctuations in the $z$-direction. A production run of 300ns in the NPT ensemble at 303 K and 1 bar was run using the Particle Mesh Ewald method [16] to compute long-range electrostatics.

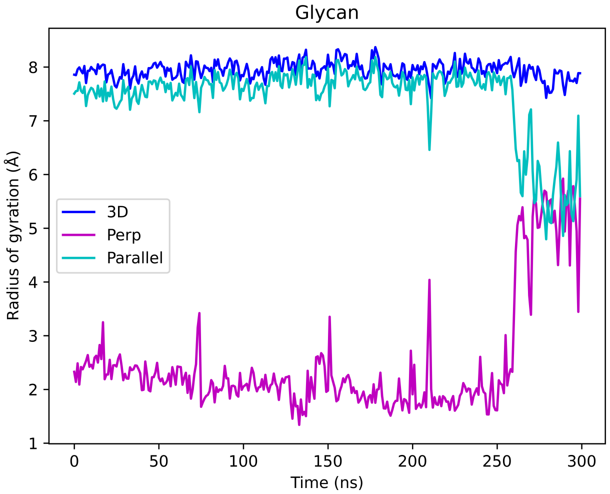


Figure S2: Parallel (cyan), perpendicular (magenta), and 3D (blue) radii of gyration of the glycan during the full simulation trajectory. Note that $R_{g3D}$ can not distinguish between a globular structure in solution and distinct configurations upon surface adsorption vs. the parallel $R_{g\parallel}$ and perpendicular $R_{g\perp}$ radii that do.

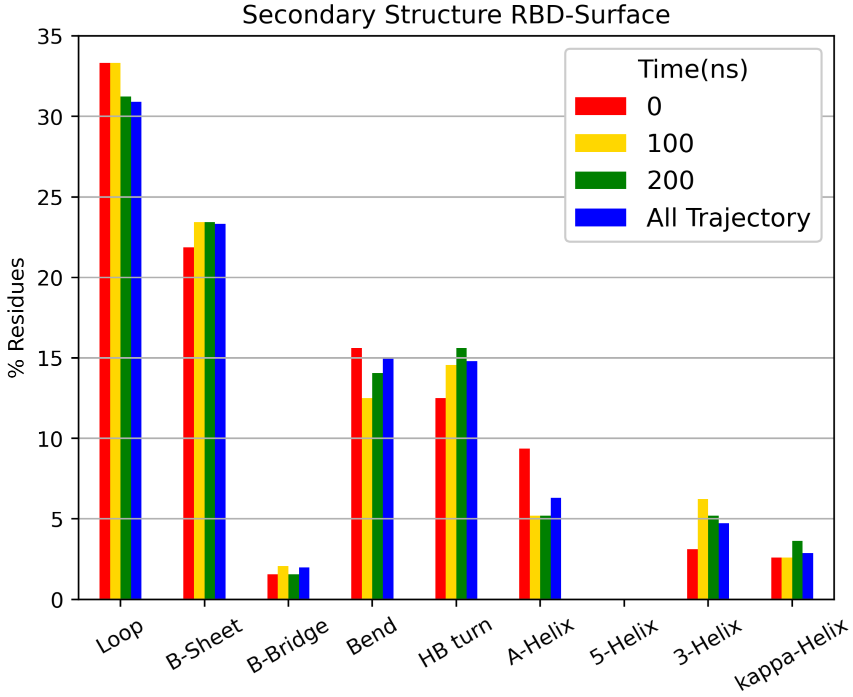


Figure S3: Secondary structure of the protein residues at the initial configuration (red), after 100ns (yellow), and after 200ns (green). The blue bars represent the mean value of residues in the respective secondary structure over the full trajectory.

| Class | Method | Property | Layer | lipid_list | Lipid | n_chain | nbins | edges | start | final | step |
|---|---|---|---|---|---|---|---|---|---|---|---|
| `Cumulative2D` | `order_matrix()` | SCD | ✓ | - | ✓ | ✓ | ✓ | ✓ | ✓ | ✓ | ✓ |
| | `all_lip_order()` | SCD | ✓ | ✓ | - | - | ✓ | ✓ | ✓ | ✓ | ✓ |
| | `height_matrix()` | Height | ✓ | ✓ | - | - | ✓ | ✓ | ✓ | ✓ | ✓ |
| | `thickness()` | Thickness | - | ✓ | - | - | ✓ | ✓ | ✓ | ✓ | ✓ |
| | `splay_matrix()` | Splay angle | ✓ | ✓ | - | - | ✓ | ✓ | ✓ | ✓ | ✓ |
| `Voronoi2D` | `voronoi_apl()` | Area per lipid | ✓ | ✓ | - | - | ✓ | ✓ | ✓ | ✓ | ✓ |
| | `voronoi_height()` | Height | ✓ | ✓ | - | - | ✓ | ✓ | ✓ | ✓ | ✓ |
| | `voronoi_thickness()` | Thickness | - | ✓ | - | - | ✓ | ✓ | ✓ | ✓ | ✓ |
| | `splay_matrix()` | Splay angle | ✓ | ✓ | - | - | ✓ | ✓ | ✓ | ✓ | ✓ |
| `PackingDefects` | `packing_defects()` | Packing Defects | ✓ | ✓ | - | - | ✓ | ✓ | ✓ | ✓ | ✓ |
| | `packing_defects_stats()` | Packing defects | ✓ | ✓ | - | - | ✓ | ✓ | ✓ | ✓ | ✓ |

Table S1: Toolbox classes and their inputs
